# Changes of urinary proteomic before and after QIV and COVID-19 vaccination

**DOI:** 10.1101/2022.03.25.485748

**Authors:** Xuanzhen Pan, Yongtao Liu, Yijin Bao, Lilong Wei, Youhe Gao

**Affiliations:** Department of Biochemistry and Molecular Biology, Beijing Key Laboratory of Gene Engineering Drug and Biotechnology, Beijing Normal University, Beijing, China 10087; Clinical Laboratory, China-Japan Friendship Hospital, Beijing, China 100029

## Abstract

We first collected a young people’s urine samples cohort of quadrivalent influenza vaccine. Urine protein at 24 hours after vaccination was enriched in immune-related pathways, though the specific pathways varied. Perhaps because different people may be in a previous life encountered some of the viruses in the vaccine, the second immunization was triggered. Or everyone has a different constitution, exposure to the same virus triggering different immunity. We then collected urine samples from several uninfected SARS-CoV-2 young people before and after the first, second, and third doses of the COVID-19 vaccine. We found that the differential protein compared between after the second dose (24h) and before the second dose enriched pathways were involved in regulated exocytosis and immune-related pathways, indicating not first exposure to antigen. Surprisingly, the urine differential protein-enriched pathways before and after the first dose were similar to those before and after the second dose. We assume that although the volunteers have not been infected with SARS-CoV-2, they might have been exposed to other coimmunogenic coronaviruses. 2~4h after the third vaccination, the differentially expressed protein also enriched regulated exocytosis and immune-related pathways, indicating that the body has triggered the immune response in a very short time after vaccination, and urine proteome is a good window to monitor the changes of human immune function.

## Introduction

Coronavirus disease 2019 (COVID-19) is an unprecedented global threat caused by severe acute respiratory syndrome coronavirus 2 (SARS-CoV-2) ^1^. About 80% of patients with COVID-19 are not severely ill, displaying mild symptoms with a good prognosis^2^. So many nations are pursuing the rollout of SARS-CoV-2 vaccines as an exit strategy from unprecedented COVID-19-related restrictions^3^. So far, however, the effectiveness of vaccines has been assessed by measuring blood indicators, and we are exploring whether urine proteins reflect changes in the body’s immunity before and after vaccination. To the best of our knowledge, there are no studies of urine proteome changes before and after vaccination.

## Materials and methods

### Urine sample from Quadrivalent influenza vaccine and COVID-19 vaccine recipients

QIV(Quadrivalent influenza vaccine) cohort of 8 volunteers (young healthy individuals) comprising 45 specimens include 6 time points. The detailed individual descriptions including age, sex and the time points of sampling are shown in **Supplementary Table1**. 6 time points contains before vaccination(T_0_); 24 hours after vaccination(T_1_); 7days(T_2_), 14days(T_3_), 21days(T_4_), 28days(T_5_) after vaccination.

COVID-19 (1) cohort of 15 volunteers (young healthy individuals) comprising 88 specimens include 7 time points. The detailed individual descriptions including age, sex and the time points of sampling are shown in **Supplementary Table1**. 7 time points contains before the first vaccination(T_0_); 24 hours after the first vaccination(T_1_); after 21days, before the second vaccination(T_2_); 24 hours after the second vaccination(T_3_); 7days(T_4_), 14days(T_5_), 21days(T_6_) after the second vaccination.

COVID-19 (2) cohort of 13 volunteers (young healthy individuals) comprising 37 specimens include 3 time points. The detailed individual descriptions including age, sex and the time points of sampling are shown in **Supplementary Table1**. 3 time points contains before the third vaccination(booster shots for COVID-19)(T_0_); first urination after vaccination(2~4 hours after vaccination)(T_1_); w7days after vaccination(T_2_).

All COVID-19 vaccine recipients tested negative for nucleic acid and had been not previously infected with SARS-CoV-2. Inactivated COVID-19 Vaccine (Vero cells), also called *CoronaVac*, was produced by SINOVAC Biotech Ltd, Beijing. This study’s ethics approval was approved by the China-Japan Friendship Hospital review boards, and each participant signed informed consent.

### Urine sample preparation for label-free analysis

After collection, the urine samples were centrifuged at 3000 ×g for 30 min at 4 °C and then stored at −80 °C. For urinary protein extraction, the urine samples were first centrifuged at 12,000 ×g for 30 min at 4 °C. Then, 15 mL of urine from each sample was precipitated with three volumes of ethanol at −20 °C overnight. The pellets were dissolved in lysis buffer (8 mol/L urea, 2 mol/L thiourea, 50 mmol/L Tris, and 25 mmol/L dithiothreitol). Finally, the supernatants were quantified by the Bradford assay. A total of 100 μg of protein was digested with trypsin (Trypsin Gold, Mass Spec Grade, Promega, Fitchburg, WI, USA) using filter-aided sample preparation (FASP) methods^4^. The protein in each sample was loaded into a 10-kDa filter device (Pall, Port Washington, NY, USA). After washing two times with urea buffer (UA, 8 mol/L urea, 0.1 mol/L Tris-HCl, pH 8.5) and 25 mmol/L NH4HCO3 solutions, the protein samples were reduced with 20 mmol/L dithiothreitol at 37 °C for 1 h and alkylated with 50 mmol/L iodoacetamide (IAA, Sigma) for 45 min in the dark. The samples were then washed with UA and NH4HCO3 and digested with trypsin (enzyme-to-protein ratio of 1:50) at 37 °C for 14 h. The digested peptides were desalted using Oasis HLB cartridges (Waters, Milford, MA, USA) and then dried by vacuum evaporation (Thermo Fisher Scientific, Bremen, Germany).

The digested peptides were dissolved in 0.1% formic acid and diluted to a concentration of 0.5 μg/μL. To generate the spectral library for DIA analysis, a pooled sample (1~2 μg of each sample) was loaded onto an equilibrated, high-pH, reversed-phase fractionation spin column (84,868, Thermo Fisher Scientific). A step gradient of 8 increasing acetonitrile concentrations (5, 7.5, 10, 12.5, 15, 17.5, 20 and 50% acetonitrile) in a volatile high-pH elution solution was then added to the columns to elute the peptides as eight different gradient fractions. The fractionated samples were then evaporated using vacuum evaporation and resuspended in 20 μL of 0.1% formic acid. Two microliters of each fraction were loaded for LC-DDA-MS/MS analysis.

### Liquid chromatography and Mass spectrometry

The iRT reagent (Biognosys, Switzerland) was added at a ratio of 1:10 v/v to all peptide samples to calibrate the retention time of the extracted peptide peaks. For analysis, 1 μg of peptide from each sample was loaded into a trap column (75 μm * 2 cm, 3 μm, C18, 100 Å) at a flow rate of 0.55 μL/min and then separated with a reversed-phase analytical column (75 μm * 250 mm, 2 μm, C18, 100 Å). Peptides were eluted with a gradient of 3%–90% buffer B (0.1% formic acid in 80% acetonitrile) for 120 min and then analyzed with an Orbitrap Fusion Lumos Tribrid Mass Spectrometer (Thermo Fisher Scientific, Waltham, MA, USA). 120 Min gradient elution: 0 min, 3% phase B; 0 min-3 min, 8% phase B; 3 min-93 min, 22% phase B; 93 min-113 min, 35% phase B; 113 min-120 min, 90% phase B. The LC settings were the same for both the DDA-MS and DIA-MS modes to maintain a stable retention time.

For the generation of the spectral library (DIA), the eight fractions obtained from the spin column separation was analyzed with mass spectrometry in DDA mode. The MS data were acquired in high-sensitivity mode. A full MS scan was acquired within a 350– 1200 m/z range with the resolution set to 120,000. The MS/MS scan was acquired in Orbitrap mode with a resolution of 30,000. The HCD collision energy was set to 30%. The AGC target was set to 4e5, and the maximum injection time was 50 ms. The individual samples were analyzed in DDA/DIA-MS mode. The variable isolation window of the DIA method with 29 windows was used for DIA acquisition(**Supplementary Table2**). The full scan was obtained at a resolution of 120,000 with a m/z range from 400 to 1200, and the DIA scan was obtained at a resolution of 30,000. The AGC target was 1e5, and the maximum injection time was 50 ms. The HCD collision energy was set to 35%.

### Mass spectrometry data processing

The Ms data of QIV cohort and COVID-19 (1) male cohort(DDA MS data) is performed label-free quantitative comparisons. Three technical replicates were injected for each sample. Base peak chromatograms were inspected visually in Xcalibur Qual Brower version 4.0.27.19(Thermo Fisher Scientific). RAW files were processed by MaxQuant version 1.6.17.0 (http://www.maxquant.org) using default parameters unless otherwise specified^5–7^. All RAW files of everyone were analyzed together in a single MaxQuant run. Database searches were performed using the Andromeda search engine included with the MaxQuant release^8^ with the Uniprot human sequence database (November 27, 2020; 196,211 sequences). Precursor mass tolerance was set to 4.5 ppm in the main search, and fragment mass tolerance was set to 20 ppm. Digestion enzyme specificity was set to Trypsin/P with a maximum of 2 missed cleavages. A minimum peptide length of 7 residues was required for identification. Up to 5 modifications per peptide were allowed; acetylation (protein N-terminal) and oxidation (Met) were set as variable modifications, and carbamidomethyl (Cys) was set as fixed modification. No Andromeda score threshold was set for unmodified peptides. A minimum Andromeda score of 40 was required for modified peptides. Peptide and protein false discovery rates (FDR) were both set to 1% based on a target-decoy reverse database. Proteins that shared all identified peptides were combined into a single protein group. If all identified peptides from one protein were a subset of identified peptides from another protein, these proteins were combined into that group. Peptides that matched multiple protein groups (“razor” peptides) were assigned to the protein group with the most unique peptides. “Match between run” based on accurate m/z and retention time was enabled with a 0.7 min match time window and 20 min alignment time window. Label-free quantitation (LFQ) was performed using the MaxLFQ algorithm built into MaxQuant^9^. Peaks were detected in Full MS, and a three-dimensional peak was constructed as a function of peak centroid m/z (7.5 ppm threshold) and peak area over time. Following de-isotoping, peptide intensities were determined by extracted ion chromatograms based on the peak area at the retention time with the maximum peak height. And peptide intensities were normalized to minimize overall proteome difference based on the assumption that most peptides do not change in intensity between samples. Protein LFQ intensity was calculated from the median of pairwise intensity ratios of peptides identified in two or more samples and adjusted to the cumulative intensity across samples. Quantification was performed using razor and unique peptides, including those modified by acetylation (protein N-terminal) and oxidation (Met). A minimum peptide ratio of 1 was required for protein intensity normalization, and “Fast LFQ” was enabled. Only proteins that were quantified by at least two unique peptides were used for analysis.

Data processing was using Perseus version 1.6.14.0 (http://www.perseus-framework.org)^10,11^. Contaminants, reverse, and protein groups identified by a single peptide were filtered from the data set. FDR was calculated as the percentage of reverse database matches out of total forward and reverse matches. Protein group LFQ intensities were log2 transformed to reduce the effect of outliers. Protein groups missing LFQ values were assigned values using imputation. Missing values were assumed to be biased toward low abundance proteins that were below the MS detection limit, referred to as “missing not at random”, an assumption that is frequently made in proteomics studies^12,13^. Imputation was performed separately for each group from a distribution with a width of 0.3 and a downshift of 1.8.

The Ms data of COVID-19 (1) female cohort and COVID-19 (2) cohort (DIA MS data) is performed label-free quantitative comparisons. To generate a spectral library, ten DDA raw files were first searched by Proteome Discoverer (version 2.1; Thermo Scientific) with SEQUEST HT against the Uniprot human sequence database (November 27, 2020; 196,211 sequences). The iRT sequence was also added to the human database. The search allowed two missed cleavage sites in trypsin digestion. Carbamidomethyl (C) was specified as the fixed modification. Oxidation (M) was specified as the variable modification. The parent ion mass tolerances were set to 10 ppm, and the fragment ion mass tolerance was set to 0.02 Da. The Q value (FDR) cutoff at the precursor and protein levels was 1%. Then, the search results were imported to Spectronaut Pulsar (Biognosys AG, Switzerland) software to generate the spectral library ^14^.

The individual acquisition DIA files were imported into Spectronaut Pulsar with default settings. The peptide retention time was calibrated according to the iRT data. Cross-run normalization was performed to calibrate the systematic variance of the LC-MS performance, and local normalization based on local regression was used ^15^. Protein inference was performed using the implemented IDPicker algorithm to generate the protein groups^16^. All results were then filtered according to a Q value less than 0.01 (corresponding to an FDR of 1%). The peptide intensity was calculated by summing the peak areas of the respective fragment ions for MS2. The protein intensity was calculated by summing the respective peptide intensity.

Self-contrasted method was used to analysis everyone data individually of all time points. The methods included two means: comparison between groups at each time point after vaccination and before vaccination respectively, as well as comparison between groups at two adjacent time points(as shown in **Figure 1**). The differential proteins were screened with the following criteria: proteins with at least two unique peptides were allowed; fold change ≥2 or ≤ 0.5; and *P* < 0.05 by Student’s *t*-test. Group differences resulting in *P* < 0.05 were identified as statistically significant. The *P*-values of group differences were also adjusted by the Benjamini and Hochberg method^17^. The differential proteins were analyzed by Gene Ontology (GO) based on biological processes(BP), cellular components(CC), and molecular functions(MF) using DAVID^18^, and biological process from WebGestalt (http://www.webgestalt.org). Protein interaction network analysis was performed using the STRING database (https://string-db.org/cgi/input.pl) and visualized by Cytoscape (V.3.7.1)^19^ and OmicsBean work-bench (http://www.omicsbean.cn).

**Figure 1.**
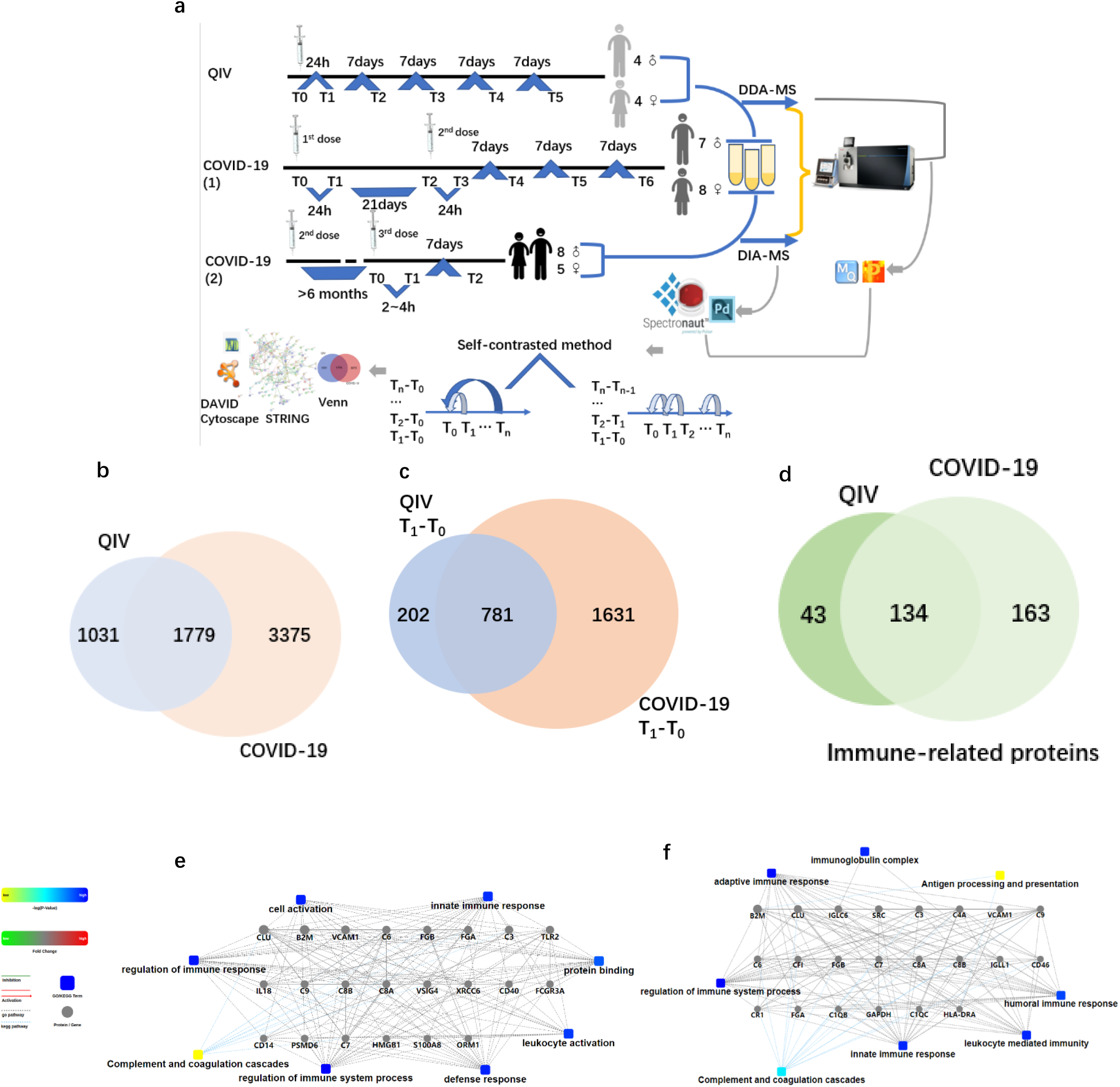
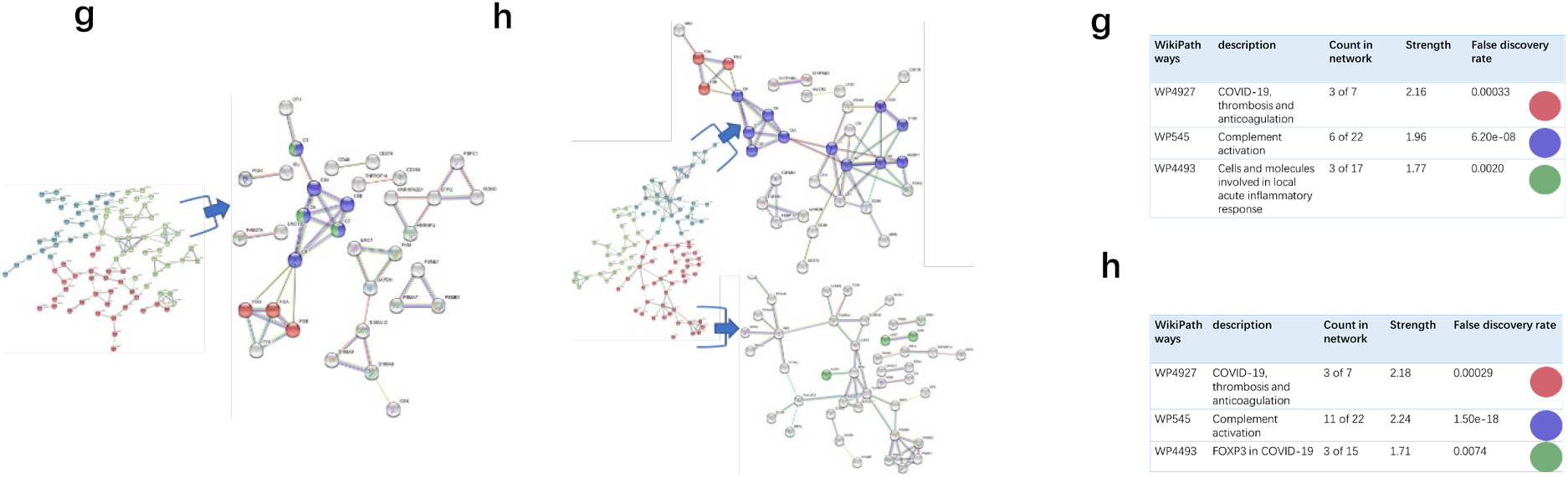
Overview of the two cohorts and the proteomic workflow. **(a)** Two cohorts and an illustration of the experimental design. A total 170 urine samples (36 vaccines) were analyzed from QIV and COVID-19 cohorts. The data-dependent/independent acquisition(DDA/DIA) technique were applied for quantitative proteomics. Integrated data analysis involved protein expression, clustering, and functional correlational network strategies. **(b)** Venn diagram of total proteins in the QIV cohort compared with the COVID-19 cohort. **(c)** Venn diagram of differential proteins in the QIV cohort compared with the COVID-19 cohort(T_1_-T_0_). **(d)** Venn diagram of immune-related proteins in the QIV cohort compared with the COVID-19 cohort. Venn diagrams showing the overlaps between total, differential(T_1_-T_0_) and immune-related proteins. **(e) and (f)** The interaction diagrams of immune-related proteins of QIV cohort and COVID-19 cohort respectively involved in tight junctions. Square box represents GO/KEGG pathways, the significance of the pathways represented by −log(*p* value) (Fisher’s exact test) was shown by color scales with dark blue as most significant. **(g)** STRING highest confidence(minimum required interaction score: 0.9) PPI network analysis of the immune-related proteins in QIV cohort. The average node degree is 1.13, average local clustering coefficient is 0.387, and PPI enrichment *p*-value is < 1.0e-16. **(h)** STRING highest confidence(minimum required interaction score: 0.9) PPI network analysis of the immune-related proteins in COVID-19 cohort. The average node degree is 1.41, average local clustering coefficient is 0.368, and PPI enrichment p-value is < 1.0e-16. The legends under illustrations of **Figure g** and **h**are on the right side of Figures, which include “count in network (The first number indicates how many proteins in the network are annotated with a particular term. The second number indicates how many proteins in total (in the network and in the background) have this term assigned)”; “strength (Log10(observed/expected).This measure describes how large the enrichment effect is. It’s the ratio between i) the number of proteins in the network that are annotated with a term and ii) the number of proteins that we expect to be annotated with this term in a random network of the same size.)”; “false discovery rate (This measure describes how significant the enrichment is. Shown are *p*-values corrected for multiple testing within each category using the Benjamini-Hochberg procedure.)”.

## Results

### Proteomic profiling of urine samples from Quadrivalent influenza vaccine and COVID-19 vaccine recipients, identification and differential proteins analysis

The QIV cohort was conducted using lable-free DDA-LC-MS/MS quantitation to characterize urinary protein profile by self-contrasted method (**Figure 1a**). A total of 2810 urinary proteins with at least 2 unique peptides were identified with <1% Q-value(corresponding to an FDR of 1%) at the protein level in all 45 samples.

The COVID-19 cohort was conducted using lable-free DDA/DIA-LC-MS/MS quantitation to characterize urinary protein profile by self-contrasted method (**Figure 1a**). A total of 5154 proteins in all samples of COVID-19 cohort; including a total of 3556 proteins in all male samples of COVID-19(1) cohort, 2061 proteins in all female samples of COVID-19(1) cohort, 1502 proteins in all samples of COVID-19(2) cohort. The comparison between the total proteins of QIV cohort and COVID-19 cohort is shown in **Figure 1b**. The comparison of the differential proteins of QIV cohort (T_1_-T_0_) and the differential protein of COVID-19 (1) cohort (T_1_-T_0_) (Fold change > 2 or < 0.5, *p*-value <0.05) is shown in **Figure 1c**. Proteins whose general function involving immune-related functions from UniProt (https://www.uniprot.org) were found(All immune-related protein were shown in **Supplementary Table 3**). There were 134 immune-related proteins in both cohorts. The comparison of immune-related proteins in the two cohorts is shown in **Figure 1d.** In addition, we also generated these immune protein-protein interactions(PPI) involved in the important biological process, KEGG pathway and molecular function respectively(**Figure 1e** and **1f**). These immune proteins contain complement C3 and so on which participate in Complement Activation. As we known complement proteins in the circulation are not activated until triggered by an encounter with a bacterial cell, a virus, an immune complex, damaged tissue or other substance not usually present in the body^20,21^. Immune proteins network is clustered to 3 sections by kmeans clustering was used in by STRING. Such as complement activation; COVID-19, thrombosis and anticoagulation; FOXP3 in COVID-19 were found in COVID-19 cohort; meanwhile, COVID-19, thrombosis and anticoagulation; complement activation and Cells and molecules involved in local acute inflammatory response were found in QIV cohort(**Figure 1g** and **1h**).

### Each influenza vaccinee’s urine proteins reflect different immune-related pathways

The urine samples of all volunteers were analyzed by self-contrasted method individually (comparison including: T_1_-T_0_, T_2_-T_0_, T_3_-T_0_, T_4_-T_0_, T_5_-T_0_; T_1_-T_0_, T_2_-T_1_, T_3_-T_2_, T_4_-T_3_, T_5_-T_4_) to obtain’ the differential proteins(fold change > 2 or < 0.5, *p-* value <0.05). The differential proteins of comparison between the time point before vaccination and the first time point after vaccination (T_1_ −T_0_) were enriched into biological processes(BP) in DAVID. We found most volunteer’s top BP contain immune-related pathways, indicating that the vaccine started working, prompting the body to initiate an immune response, although the specific immune-related pathways involved were different. The differential proteins from each person’s comparison between the other two time points were also analyzed separately for enrichment(information about BP,MF,KEGG in DVAID was shown in **Supplementary Table 4**). Total immune-related BP(*p*-value < 0.05) of everyone was shown in **Figure 2a**. The triggered immune pathways of most people were innate immune response, viral entry into host cell, acute-phase response, antibacterial humoral response, immune response, inflammatory response, leukocyte migration, receptor-mediated endocytosis. The immune-related pathways(*p*-value < 0.05) obtained from the comparison of each person at two time points were shown in the **Supplementary Table 5**. The FDR value of the unmarked pathway is less than 0.05, the FDR value of the pathway marked with light blue is less than 0.5, and the FDR value of the pathway marked with light blue is less than 0.5. Venn diagrams was used to show the overlaps between significantly changed proteins (fold change > 2 or < 0.5, *p*-value <0.05) in T_1_ compared T_0_. We did not find any common proteins, and the number of unique proteins of each person ranged from 13 to 92(**Supplementary Table 5**). These significantly regulated proteins were analyzed by omicsbean, up/down-regulate and PPI were shown in **Figure 2b**. We found that the proportion of up-regulated proteins was different in everyone at the first time point after vaccination, in addition to the different immune pathways induced. Some people involved in the immune response of the upregulation protein accounted for immune system activation; other people have more down-regulated proteins, which suppress the immune system slightly. Proteins involved in immune pathways are down-regulated. And the changed proteins involved in leukocyte transendothelial migration were all down-regulated. **Figure 2c** shows the overlap of the differential changed proteins obtained by the two comparison methods for each vaccinator. We combined the differential proteins obtained by comparison at these different time points and their fold change to obtain the overall immune system profile of each person.

**Figure 2.**
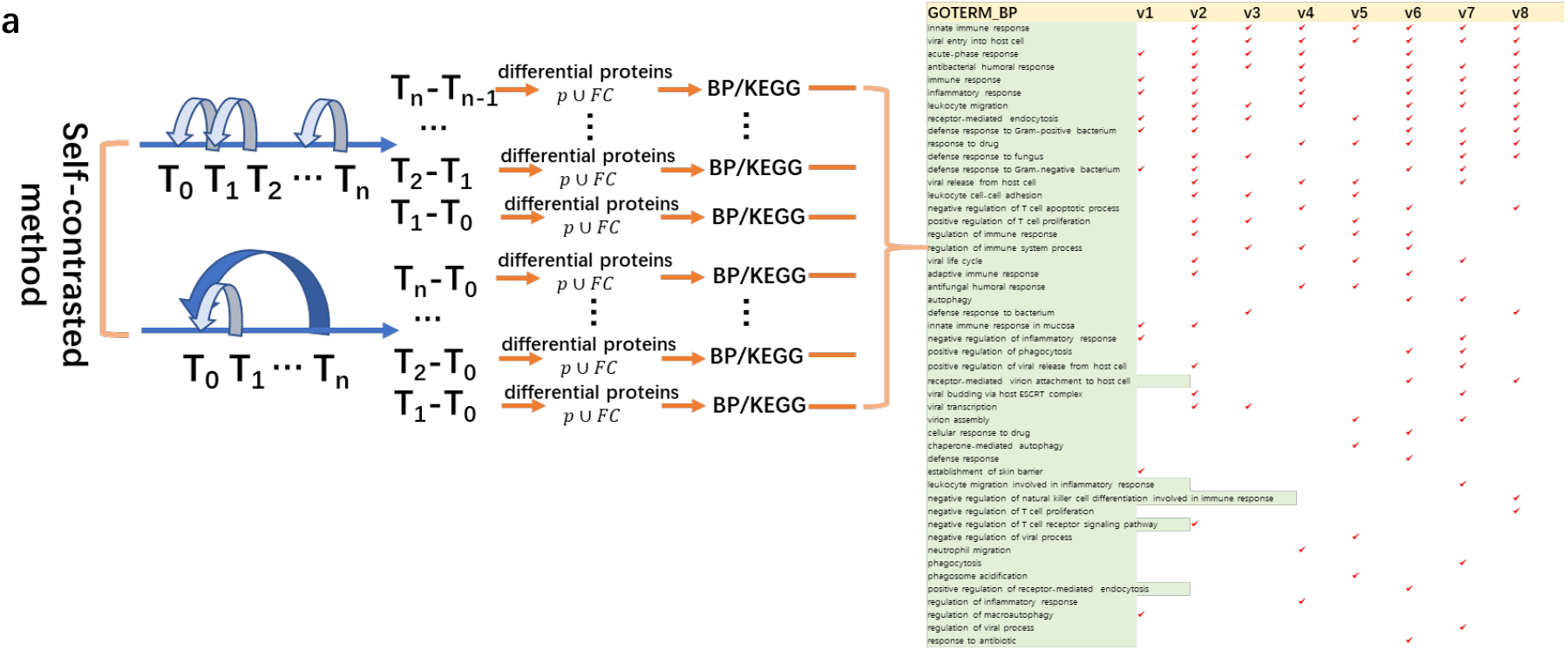
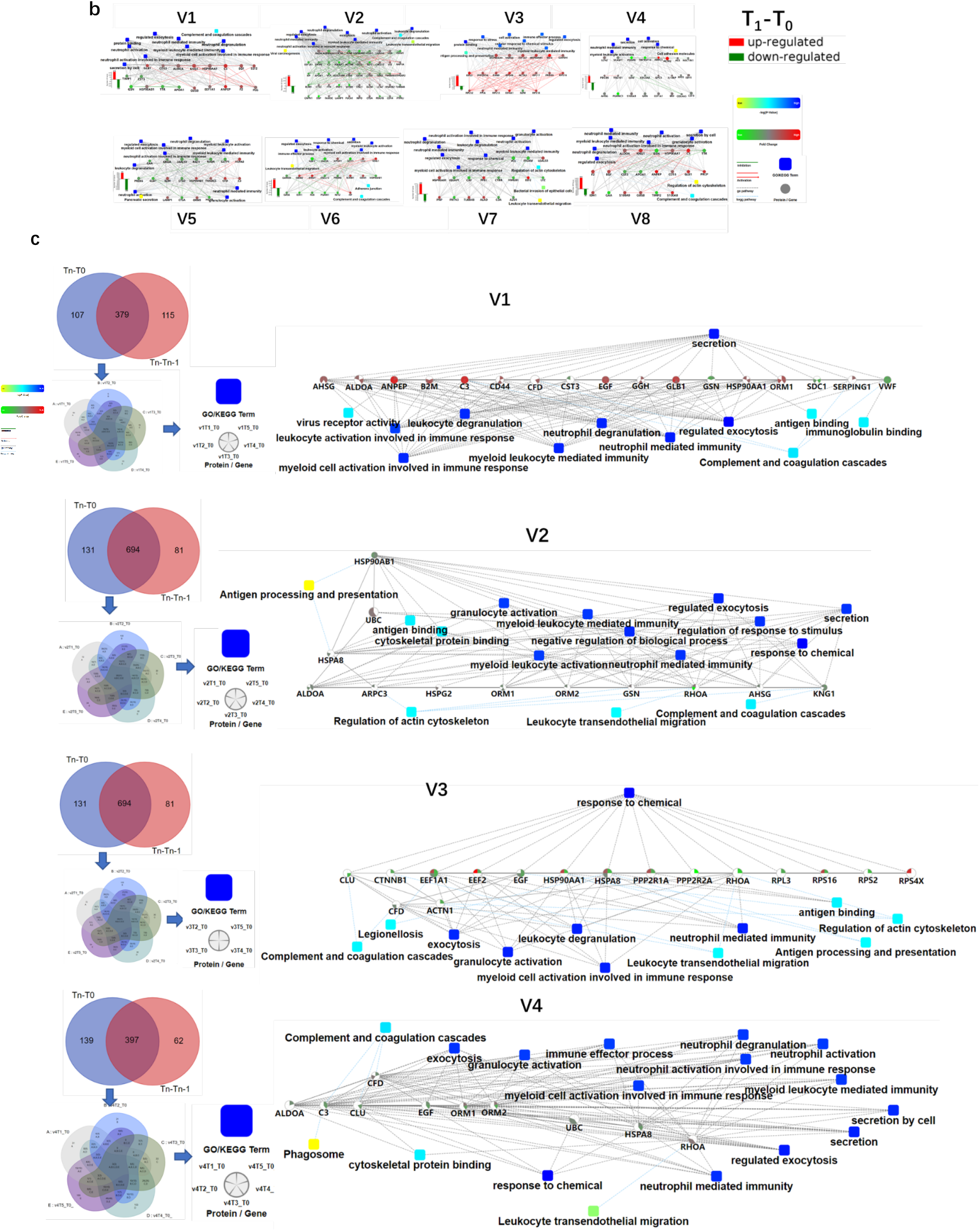
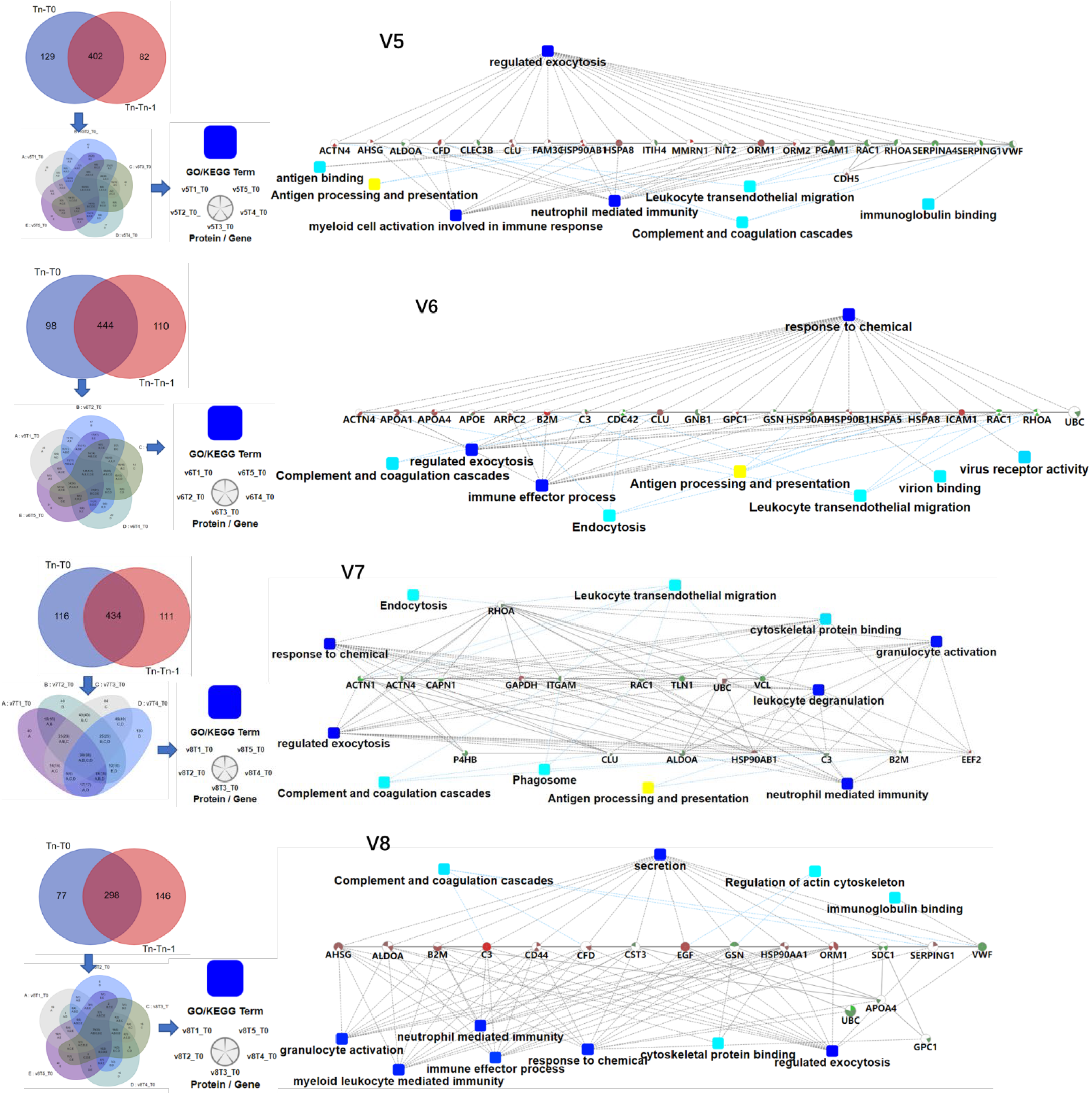
Differences in the immune response of each volunteer after QIV vaccination. **(a)** Total immune-related BP (p-value < 0.05) of everyone by DAVID. The differential proteins(fold change > 2 or < 0.5, p-value <0.05) were obtained by self-contrasted method individually (comparison including: T1-T0, T2-T0, T3-T0, T4-T0, T5-T0; T1-T0, T2-T1, T3-T2, T4-T3, T5-T4). **(b)** Significantly changed proteins (fold change > 2 or < 0.5, p-value <0.05) in T1 compared T0 was enriched by omicsbean, and their up/down-regulate and PPI were different. Network nodes and edges represent proteins and protein–protein associations. Green/red solid lines represent inhibition/activation; gray dotted lines represent GO pathways. Color bar from red to green represents the fold change of protein level from increasing to decreasing. The significance of the pathways represented by −log(p value) (Fisher’s exact test) was shown by color scales with dark blue as most significant. (**c**) Venn diagram of the significantly changed proteins (fold change > 2 or < 0.5, p-value <0.05) obtained by the two comparison methods at all different time points for each vaccinator. The proteins and their fold change to obtain the overall immune system profile of each person by STRING and omicsbean. These pathway p-value were adjusted.

### The immune-related pathways induced by the first and second doses of COVID-19 vaccine were similar, and vaccinees may had been exposed to other coronaviruses before vaccination

The urine of the COVID-19 vaccinees was collected from March 2021. None of the volunteers had previously been infected with SARS-CoV-2, and the nucleic acid test was negative. We first obtained total of 1125 differential proteins by comparing the data of each vaccinee (fold change > 2 or < 0.5, p-value <0.05) before and after the first dose of COVID-19 vaccination (T_1_-T_0_). The differentially expressed proteins were used to enrich biological process (the cut-off of p-value adjusted is set to 0.01), we found that everyone had similar enriched biological process (**Figure 3a**), including regulated exocytosis, response to chemical/stress, and immune-related pathways (belong to top 40 BP, p-value adjusted < e-10). The pathway activation strength value of enriched biological process was shown in **Figure 3b**, suggesting that the immune responses are suppressed, activated or concurrence. All vaccinees’ differentially expressed proteins were simultaneously processed by omicsbean for gene ontology analysis, levels of enriched biological process, KEGG pathway and molecular function were shown in **Figure 3c**. Next, we obtained total of 1461 differential proteins by comparing the data of each vaccinee (fold change > 2 or < 0.5, p-value <0.05) before and after the second dose of COVID-19 vaccination (T_3_-T_2_). The differentially expressed proteins were used to enrich biological process (the cut-off of p-value adjusted is set to 0.01), we found enriched biological process (**Figure 3d**)also included regulated exocytosis, response to chemical/stress/stimulus, and immune-related pathways (belong to top 40 BP, p-value adjusted < e-5).

**Figure 3.**
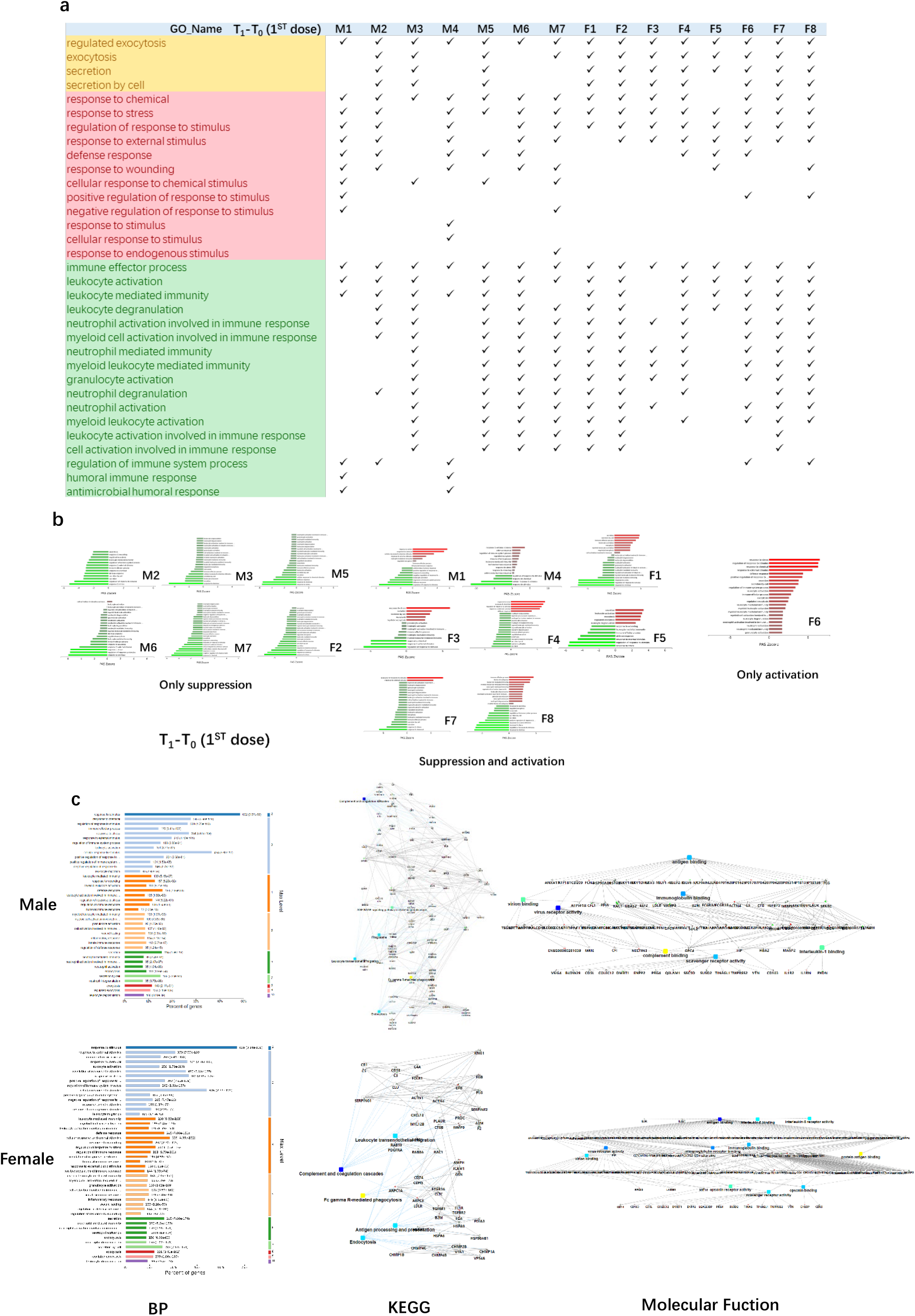
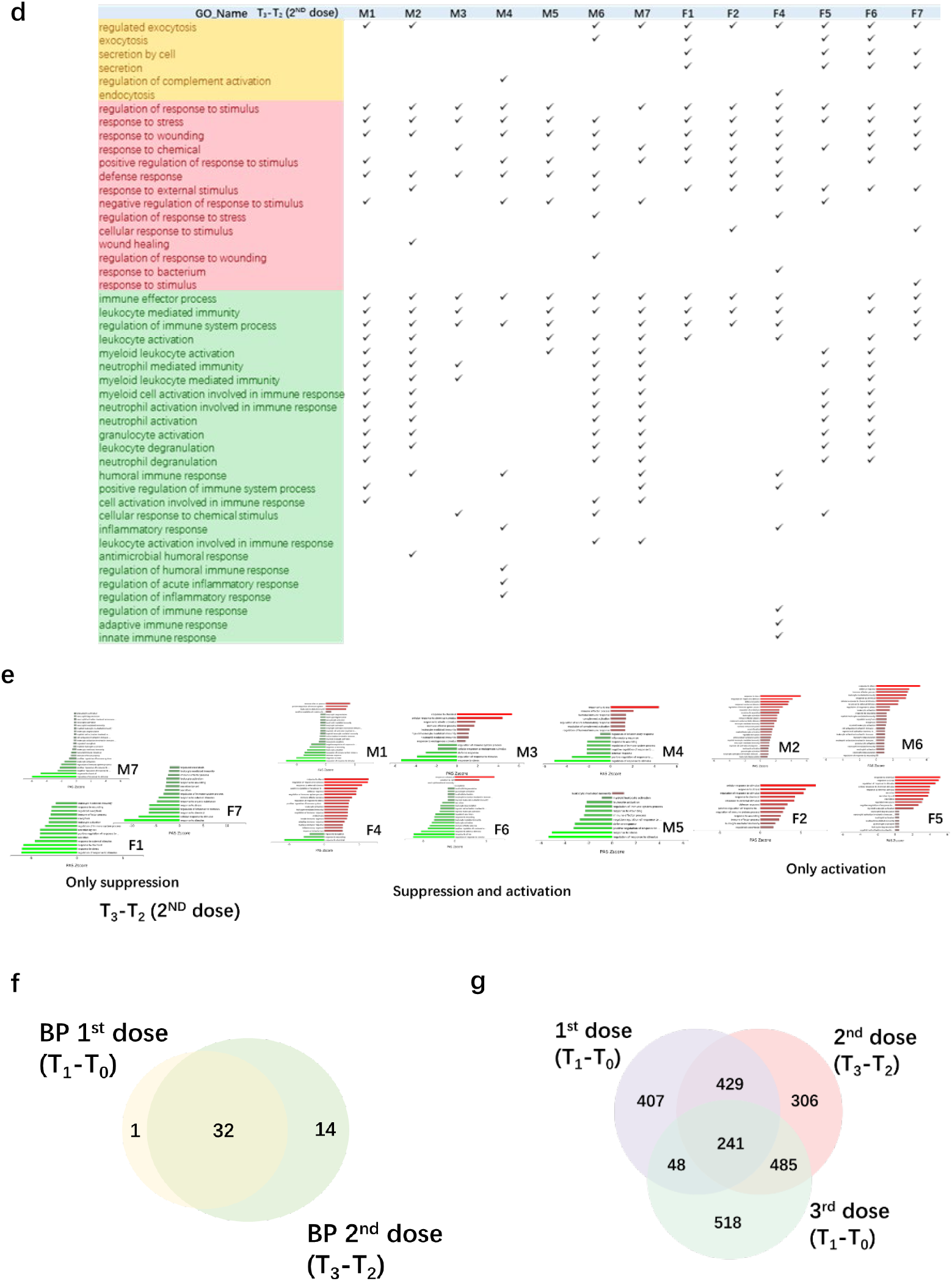
Functional analysis of differentially changed proteins in the first and second dose of COVID-19 vaccinees. **(a)** Demonstration of enriched top 40 immune-related biological processes by differential proteins(fold change > 2 or < 0.5, p-value <0.05) in the comparison between T_0_(before the first dose of vaccination) and T_1_(24h after the vaccination) of each vaccine, p-value adjusted < e-10. **(b)** The pathway activation strength value of enriched immune-related biological process(T_1_-T_0_), which served as the activation profiles of the Signaling pathways based on the expression of individual genes. Vaccinees immune responses were suppressed, activated or concurrence. The z-score algorithm was used to predict the activation state (either activated or inhibited)30 of the biological process. If the z-score ≤ −2, the process is predicted to be statistically significantly inhibited. (**c**) GO and KEGG pathway were enriched for this dataset (T_1_-T_0_), these enriched processes are statistically significant with p-value adjusted(p-value: calculated with Fish exact test with Hypergeometric algorithm; p-value adjusted: using ‘Benjamini-Hochberg’ method for multiple tests). Network nodes and edges represent proteins and protein–protein associations. Green/red solid lines represent inhibition/activation; gray dotted lines represent GO pathways. Color bar from red to green represents the fold change of protein level from increasing to decreasing. The significance of the pathways represented by −log(p value) (Fisher’s exact test) was shown by color scales with dark blue as most significant. (**d**) Demonstration of enriched top 40 immune-related biological processes by differential proteins(fold change > 2 or < 0.5, p-value <0.05) in the comparison between T_2_(before the second dose of vaccination) and T_3_(24h after the vaccination) of each vaccine, p-value adjusted < e-5. **(b)** The pathway activation strength value of enriched immune-related biological process(T_3_-T_2_). More individuals with upregulated immune systems. **(f)** Venn diagram of significant top 40 immune-related pathways showing the overlaps between the first and second doses of COVID-19 vaccine(T_1_-T_0_; T_3_-T_2_). **(g)** Venn diagram showing the overlaps of total differentially expressed proteins among before and after 1st dose, 2nd dose, and 3rd dose(T_1_-T_0_^1st^; T_3_-T_2_^2nd^; T_1_-T_0_^3rd)^.

The pathway activation strength value of enriched biological process was shown in **Figure 3e**, suggesting that most vaccinees’ immune systems were activated after the second dose, compared to the pathways enriched after the first dose. The immune-related pathways overlap between the first and second doses of COVID-19 vaccine were similar(**Figure 3f**), most biological processes of the immune-related pathways enriched by the first dose were included in second dose. The venn diagrams showed total differentially expressed proteins overlap among before and after 1st dose, 2nd dose, and 3rd dose(**Figure 3g**).

### Immune-related pathways can be enriched when vaccinees urinate for the first time(2~4h) after the third dose of COVID-19 vaccine

We collected urine samples from vaccinees who accepted the third (booster) dose COVID-19 vaccine in November 2021, 6 to 9 months after the first and second dose vaccination(**Figure 4a**). Before the vaccination(T_0_), vaccinees’ urine samples were collected. Within 2 ~ 4 h after vaccination(T_1_), the first urine samples excreted by the vaccinees after vaccination were collected. The last urine samples(T_2_) were collected after a week(7 days). At the same time, we also set up a control sample, the volunteer did not accept the third dose of vaccine. We collected his urine samples at two time points(T_0_ and T_2_).

**Figure 4.**
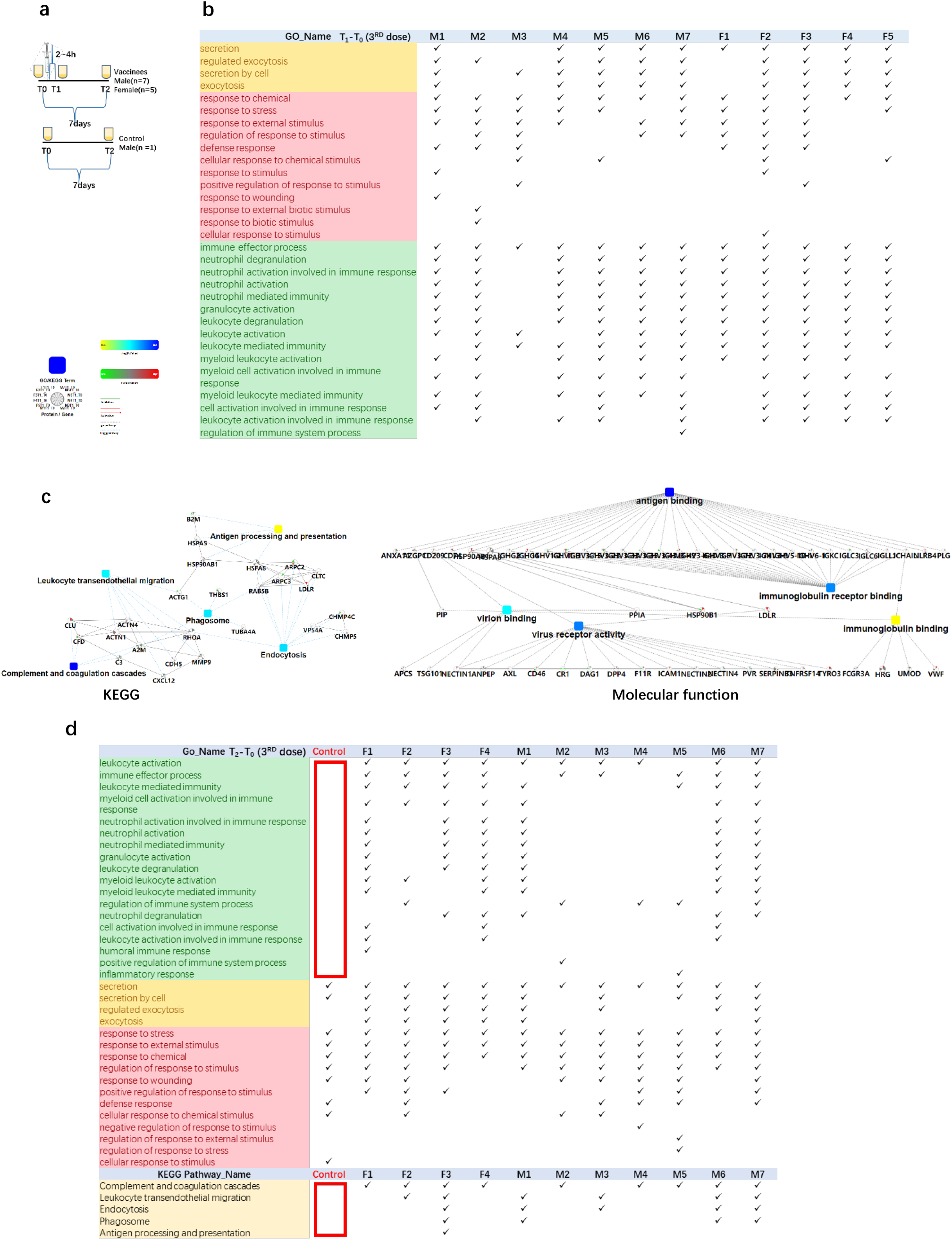
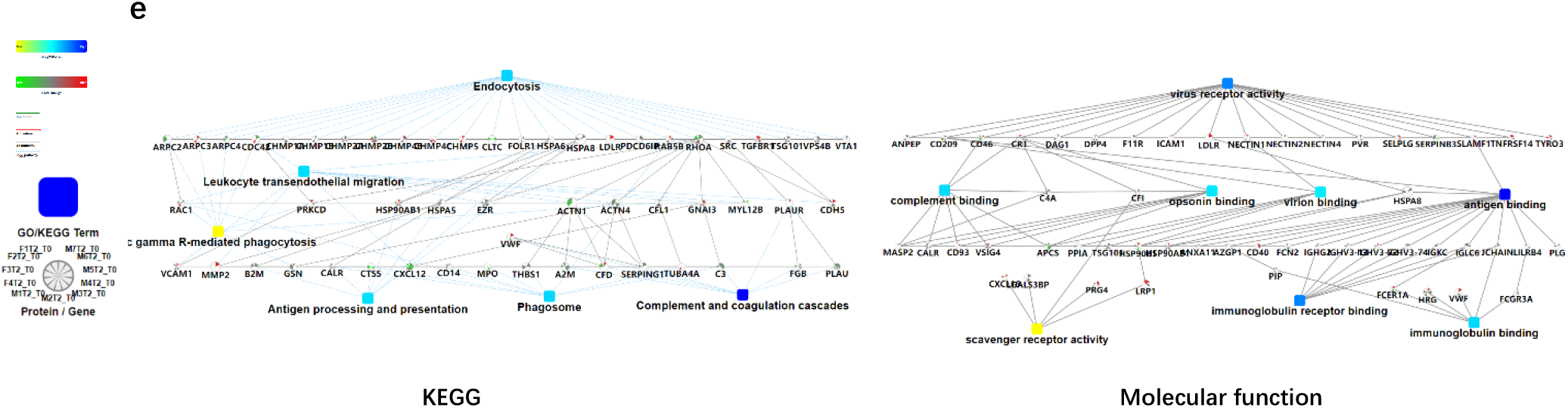
Functional analysis of differentially changed proteins in the third dose(booster dose) of COVID-19 vaccinees. **(a)** Design of the times point of sampling between the vaccinees and the control. All of them had received the first and the second doses of vaccine for more than six months. **(b)** Demonstration of enriched top 40 immune-related biological processes by differential proteins(fold change > 2 or < 0.5, p-value <0.05) in the comparison between T_0_(before the third dose of vaccination) and T_1_(2~4h after the vaccination, the first urination after vaccination) of each vaccine, p-value adjusted < e-12. **(c)** The interaction diagrams showing significant pathways including KEGG pathway and molecular function (T_1_-T_0_^3rd^). The KEGG pathways contains complement and coagulations cascades, endocytosis, phagosome, leukocyte transendothelial migration and antigen processing and presentation(p-value adjusted < 1.1e-2). The enriched molecular function contains antigen binding, virus receptor activity, virion binding, immunoglobulin receptor binding(p-value adjusted < 5.70e-23). (**d**) Demonstration of enriched top 40 significant immune-related biological processes and KEGG pathways (p-value adjusted < 0.05)by differential proteins(fold change > 2 or < 0.5, p-value <0.05) in the comparison between T_0_(before the third dose of vaccination) and T_2_(7days after the vaccination) of each vaccine. **(e)** The interaction diagrams(T_2_-T_0_^3rd^) showing significant pathways including KEGG pathway(p-value adjusted < 1.18e-3) and molecular function(p-value adjusted < 5.35e-4). (p-value adjusted < 1.18e-3). Network nodes and edges represent proteins and protein-protein associations. Green/red solid lines represent inhibition/activation; gray dotted lines represent GO pathways. Color bar from red to green represents the fold change of protein level from increasing to decreasing. The significance of the pathways represented by −log(p value) (Fisher’s exact test) was shown by color scales with dark blue as most significant. P-value adjusted was used ‘Benjamini-Hochberg’ method for multiple tests

The immune response could be reflected in the first urination after vaccination. Total of 1292 differential changed proteins (fold change > 2 or < 0.5, p-value <0.05)enriched in secretion/regulated exocytosis, response to chemical/stress/stimulus, and immune-related pathways. The immune-related biological processes of top 40 were in **Figure 4b**(p-value adjusted < e-12). The KEGG pathways contains complement and coagulations cascades, endocytosis, phagosome, leukocyte transendothelial migration and antigen processing and presentation(p-value adjusted < 1.1e-2). The enriched molecular function contains antigen binding, virus receptor activity, virion binding, immunoglobulin receptor binding(p-value adjusted < 5.70e-23)(**Figure 4c**).

Next, we found that the differential changed proteins in the control sample before and after one week only contained secretion(T_2_-T_0_), response to stimulus and other pathways, but did not contain immune-related pathways(**Figure 4d**). KEGG pathways(p-value adjusted < 0.05) such as complement and coagulation cascades, endocytosis, phagosome, leukocyte transendothelial migration and antigen processing and presentation were also not involved(**Figure 4d**). On the contrary, the immune-related pathways were enriched in the differential changed proteins comparison T2 and T0 time point of vaccinees urine samples individually(**Figure 4d**). The KEGG pathways contains complement and coagulations cascades, endocytosis, phagosome, leukocyte transendothelial migration and antigen processing and presentation(p-value adjusted < 1.18e-3). The enriched molecular function contains virus receptor activity, antigen binding, virion binding, complement binding, immunoglobulin binding, immunoglobulin receptor binding, scavenger receptor activity(p-value adjusted < 5.35e-4)(**Figure 4e**).

## Discussion

For the first time, we explored the immune response process after vaccination from the perspective of urine proteome. We found that even after vaccination with the same vaccine, the differentially expressed proteins in the urine proteome were enriched into different immune-related pathways. Our exploration may provide a new idea for vaccine validation in the future, which can verify the efficacy of vaccine relatively earlier. As for reasons of each person’s immune response varying after the same quadrivalent influenza vaccine, we assume that because different people may be in a previous life encounter the four kinds of virus, some people may be more or less contact, the second immunization was triggered. Or we could speculate that different people have different constitutions, so exposure to the same virus triggers different levels of immunity.

Then COVID-19 outbreaks giving us inspiration, we collected a batch of COVID-19 vaccine urine samples of young people, because these volunteers had not been infected with the SARS-CoV-2, and negative for nucleic acid test. It’s certain to say they haven’t been exposed to the virus before. Perhaps in this case, the volunteers’ urine protein caused by immune response after vaccination should be close to or unified. However, it turned out that the differential expressed proteins and the specific immune response pathways were different from person to person. We found that the biological processes involved by the differential changed proteins before and after the first vaccination were similar to those caused by the differential changed proteins before and after the second vaccination, including response to stimulus, secretion and other immune-related pathways.

Thus, we speculated that volunteers before vaccination may contact other kinds of coronavirus, who had triggered the primary immune response previously, and the first vaccination is equivalent to the secondary immune response. So, the pathways are analogous to the second vaccination. Studies have shown that the mortality rate and severe illness rate of those who received the first two doses of COVID-19 vaccine after re-exposure to the mutated SARS-CoV-2 virus have decreased, which is consistent with our results, and also confirms that part of the early cases of COVID-19 patients were only mild, which is probably due to their exposure to other coronavirus before exposure to the SARS-CoV-2. In the elderly, mortality and severe illness may be increased due to decreased immune response^22,23^. Finally, we collected volunteers’ urine from their first excretion after the third vaccination, and surprisingly, the immune response could be reflected in the urinary protein only 2~4 hours later.

In conclusion, we found that urinary protein have obvious change before and after vaccination, and the significant proteins belong to regulated exocytosis and immune response etc. Urinary protein could reveal the body’s immune response, to provide new ideas for the vaccine efficacy test. Different people were found to have different immune response mechanisms triggered by the same vaccine. It also confirmed the role and necessity of COVID-19 Vaccine.

## Author contributions

X. P. designed and conducted the experiments, analyzed the data, prepared the figures, and wrote the paper. Y.L. and Y. B provided guidance and help. L.W collected experimental samples. Y.G. provided the main ideas, designed experiments, and guided writing the paper.

## Data Availability Statement

The proteomic raw data are available in iProX Datasets.

